# Identification of 2019-nCoV related coronaviruses in Malayan pangolins in southern China

**DOI:** 10.1101/2020.02.13.945485

**Authors:** Tommy Tsan-Yuk Lam, Marcus Ho-Hin Shum, Hua-Chen Zhu, Yi-Gang Tong, Xue-Bing Ni, Yun-Shi Liao, Wei Wei, William Yiu-Man Cheung, Wen-Juan Li, Lian-Feng Li, Gabriel M Leung, Edward C. Holmes, Yan-Ling Hu, Yi Guan

**Affiliations:** Joint Institute of Virology (Shantou University / The University of Hong Kong) & Guangdong-Hongkong Joint Laboratory of Emerging Infectious Diseases, Shantou University, Shantou, Guangdong, 515063, P. R. China; State Key Laboratory of Emerging Infectious Diseases, School of Public Health, The University of Hong Kong, Hong Kong SAR, P. R. China; Beijing Advanced Innovation Center for Soft Matter Science and Engineering (BAIC-SM), College of Life Science and Technology, Beijing University of Chemical Technology, Beijing, 100029, P. R. China; Life Sciences Institute, Guangxi Medical University, Nanning, Guangxi, 530021, P. R. China; Marie Bashir Institute for Infectious Diseases and Biosecurity, School of Life and Environmental Sciences and School of Medical Sciences, The University of Sydney, Sydney, Australia

## Abstract

The ongoing outbreak of viral pneumonia in China and beyond is associated with a novel coronavirus, provisionally termed 2019-nCoV. This outbreak has been tentatively associated with a seafood market in Wuhan, China, where the sale of wild animals may be the source of zoonotic infection. Although bats are likely reservoir hosts for 2019-nCoV, the identity of any intermediate host facilitating transfer to humans is unknown. Here, we report the identification of 2019-nCoV related coronaviruses in pangolins (*Manis javanica*) seized in anti-smuggling operations in southern China. Metagenomic sequencing identified pangolin associated CoVs that belong to two sub-lineages of 2019-nCoV related coronaviruses, including one very closely related to 2019-nCoV in the receptor-binding domain. The discovery of multiple lineages of pangolin coronavirus and their similarity to 2019-nCoV suggests that pangolins should be considered as possible intermediate hosts for this novel human virus and should be removed from wet markets to prevent zoonotic transmission.

## Text

An outbreak of serious pneumonia disease was reported in Wuhan, China on 30 December 2019. The causative agent was soon identified as a novel coronavirus - 2019-nCoV^1^. Case numbers grew rapidly from 27 in December 2019 to 7,818 globally as of 30 January 2020^2^, leading to the WHO to declare a public health emergency. Many of the early cases were linked to Huanan seafood market in Wuhan city, Hubei province, from where the probable zoonotic source is speculated to originate^3^. Currently, only environmental samples taken from the market have been reported as positive for 2019-nCoV by the China CDC^4^. However, as similar wet markets were already implicated in the SARS outbreak of 2002-2003^5^, it seems likely that wild animals are also involved in the emergence of 2019-nCoV. Indeed, a number of non-aquatic mammals were available for purchase in the Huanan seafood market prior to the outbreak^4^. Unfortunately, because the market was cleared soon after the outbreak began, determining the source virus in the wild animals from the market is challenging. Although a coronavirus closely related to 2019-nCoV sampled from a *Rhinolophus affinis* bat in Yunnan in 2013 has now been identified^6^, closely related viruses have not yet been detected in other wildlife species. Here, we present the identification of 2019-nCoV related viruses in pangolins smuggled into southern China.

We investigated the virome composition of pangolins (mammalian order Pholidota). These animals are of growing importance and interest because they are the most illegally trafficked of any group of mammal: they are used as both a food source and their scales are utilized in traditional Chinese medicine. A number of pangolin species now are regarded as critically endangered on the International Union for Conservation of Nature Red List of Threatened Species. We received frozen tissue (lungs, intestine, blood) samples that were collected from 18 Malayan pangolins (*Manis javanica*) during August 2017-January 2018. These pangolins were obtained during the anti-smuggling operations by Guangxi Customs. Strikingly, high-throughput sequencing of their RNA revealed the presence of coronaviruses in six (two lung, two intestine, one lung-intestine mix, one blood) of 43 samples. With the sequence read data, and by filling gaps with amplicon sequencing, we were able to obtain six full or nearly full genome sequences - denoted GX/P1E, GX/P2V, GX/P3B, GX/P4L, GX/P5E and GX/P5L - that fall into the 2019-nCoV lineage (within the genus *Betacoronavirus*) in a phylogenetic analysis (Figure 1a). These viruses also have similar genomic organization as 2019-nCoV, with nine predicted open reading frames (Figure 1b; Extended Data Table S5). We were also able to successfully isolate the virus using the Vero E6 cell line (Extended Data Figure S1).

**Figure 1.**
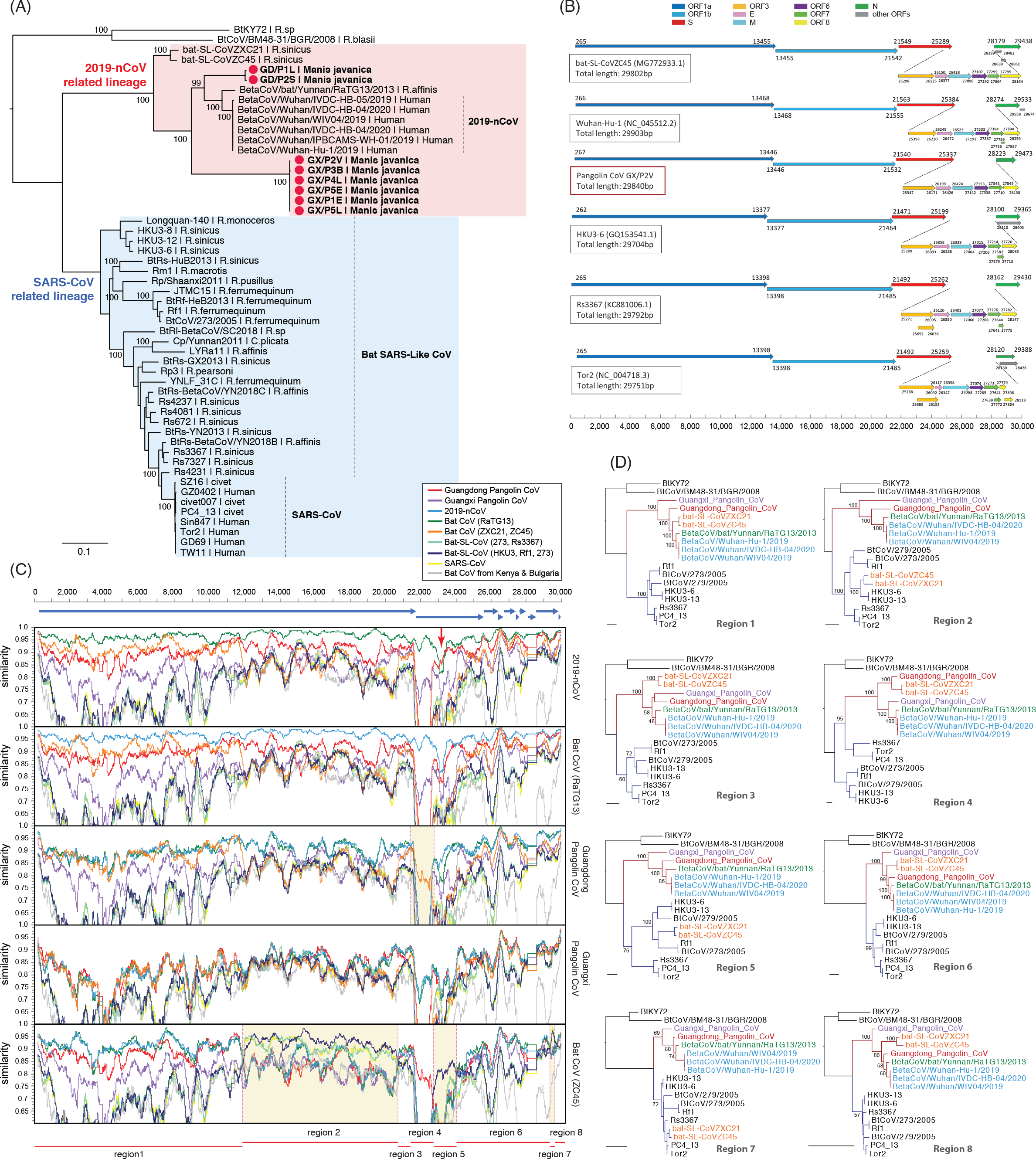
Phylogenetic analyses depicting the evolutionary relationship between human 2019-nCoV, the pangolin coronavirus sequences obtained in this study, and the other reference coronaviruses. The phylogenies were estimated using a maximum likelihood approach employing the GTR+I+Γ nucleotide substitution model and 1,000 bootstrap replicates. (A) Phylogeny of the subgenus *Sarbecovirus* (genus *Betacoronavirus*) estimated from the concatenated ORF1ab-S-E-M-N genes. Red circles indicate the pangolin coronavirus sequences generated in this study. Note that GD/P1L is the consensus sequence re-assembled from the raw data previously published^7^. (B) Genome organization of coronaviruses including the pangolin coronaviruses, with the predicted ORFs shown in different colors. (C) Sliding window analysis of changing patterns of sequence similarity between 2019-nCoV, pangolin coronaviruses and bat coronavirus RaTG13. The name of the query sequences are shown vertically on the right of the analysis boxes. The similarities to different reference sequences are indicated by different colors shown in the legend box at the top. Guangdong pangolin coronaviruses GD/P1L and GD/P2S were merged for this analysis. The blue arrows at the top indicate the position of the ORFs in the alignment analyzed. The potential recombination breakpoints are shown in pink dash lines, which together slice the genomes into eight regions (regions with <200bp were omitted; indicated by the red line at bottom) for phylogenetic analysis. (D) Phylogenetic trees of different genomic regions. SARS-CoV and 2019-nCoV related lineages are shown in blue and red tree branches. Branch scale bars are 0.1 substitutions/site.

Based on these new genome sequences, we designed primers for qPCR detection to confirm that the raw samples were positive for the coronavirus. We conducted further qPCR testing on another batch of archived pangolin samples collected between May-July 2018. Among the 19 samples (nine intestine tissues, ten lung tissues) tested from 12 animals, three lung tissue samples were coronavirus positive.

In addition to the animals from Guangxi, after the start of the 2019-nCoV outbreak the Guangzhou Customs Technology Center re-examined their five archived pangolin samples (two skin swabs, one unknown tissue, one scale) obtained in previous anti-smuggling operations in March 2019. Following high-throughput sequencing the scale sample was found to contain coronavirus reads, and from these data we were able to assemble a partial genome sequence of 21,505bp (denoted as GD/P2S), representing approximately 72% of the 2019-nCoV genome. This virus sequence on the scale may in fact come from contaminants of other infected pangolin tissues. Notably, another study of diseased pangolins in Guangdong performed in 2019 also identified viral contigs from lung samples that were similarly related to 2019-nCoV^7^. Different assembly methods and manual curation were performed to generate a partial genome sequence that comprised about 86.3% of the full-length virus genome (denoted as GD/P1L in the Figure 1a tree).

These novel pangolin coronavirus genomes have approximately 85.5% to 92.4% similarity to 2019-nCoV, and represent two sub-lineages of 2019-nCoVs in the phylogenetic tree, one of which (GD/P1L and GDP2S) is extremely closely related to 2019-nCoV (Figure 1; red circles). It has previously been noted that members of the subgenus *Sarbecovirus* have experienced widespread recombination^8^. In support of this, a recombination analysis performed here revealed that bat coronaviruses ZC45 and ZCS21 are likely recombinants, containing genome fragments from multiple SARS-CoV related lineages (genome regions 2, 5, 7) and 2019-nCoV related lineages including that from pangolins (regions 1, 3, 4, 6, 8).

More notable, however, was the observation of putative recombination signals between the pangolins coronaviruses, bat coronaviruses RaTG13, and human 2019-nCoV (Figure 1c, d). In particular, 2019-nCoV exhibits very high sequence similarity to the Guangdong pangolin coronaviruses in the receptor-binding domain (RBD; 97.4% amino acid similarity; indicated by red arrow in Figure 1c and Figure 2a), even though it is most closely related to bat coronavirus RaTG13 in the remainder of the viral genome. Bat CoV RaTG and the human 2019-nCoV have only 89.2% amino acid similarity in RBD. Indeed, the Guangdong pangolin coronaviruses and 2019-nCoV possess identical amino acids at the five critical residues of the RBD, whereas RaTG13 only shares one amino acid with 2019-nCoV (residue 442, human SARS-CoV numbering^9^). Interestingly, a phylogenetic analysis of synonymous sites alone in the RBD revealed that the phylogenetic position of the Guangdong pangolin is consistent with that in the remainder of the viral genome, rather than being the closest relative of 2019-nCoV (Figure 2b). Hence, it is possible that the amino acid similarity between the RBD of the Guangdong pangolin coronaviruses and 2019-nCoV is due to selectively-mediated convergent evolution rather than recombination, although it is difficult to choose between these scenarios on current data. Although the drivers of any convergent evolution are unknown, its possible occurrence, as well as that of recombination, would further highlight the role played by intermediate animal hosts in human virus emergence.

**Figure 2.**
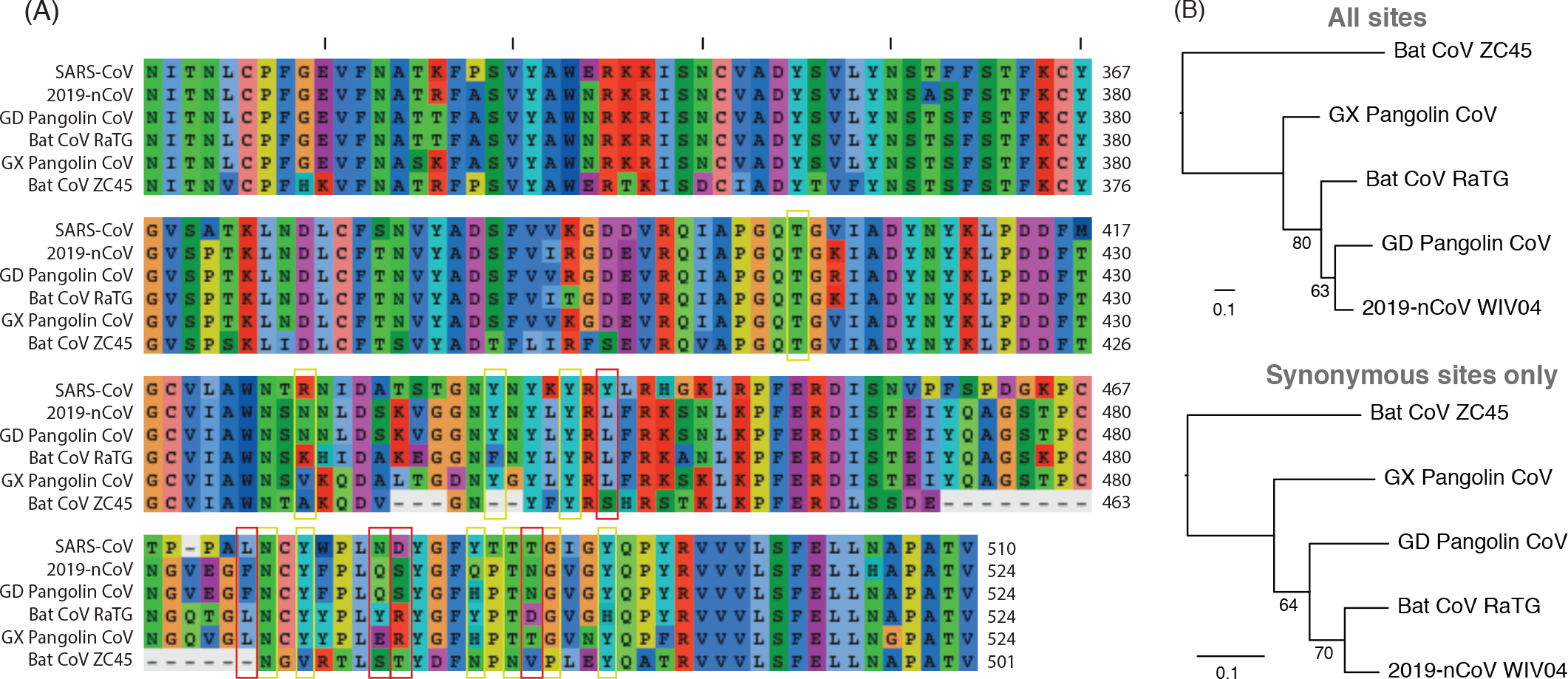
(A) Sequence alignment showing the receptor-binding domain (RBD) in human, pangolin and bat coronaviruses. The five critical residues for binding between SARS-CoV RBD and human ACE2 protein are indicated in red boxes, and ACE2-contacting residues are indicated with yellow boxes, following Wan et al.^9^. Note that in Guangdong pangolin sequence, the codon positions coding for amino acid 337 proline, 420 aspartic acid, 499 proline and 519 asparagine have ambiguous nucleotide compositions, resulting to possibly alternative amino acids threonine, glycine, threonine and lysine respectively. GD: Guangdong, GX: Guangxi. (B) Phylogenetic trees of 2019-nCoV related lineage estimated from the whole RBD region (upper) and synonymous sites only (lower). Branch supports obtained from 1,000 bootstrap replicates are shown.

To date, pangolins are the only mammals other than bats documented to be infected by a 2019-nCoV related coronavirus. It is striking that two related lineages of CoVs are found in pangolins and that both are also related to 2019-nCoV. This suggests that these animals may be long-term reservoir hosts for these viruses, which is surprising as pangolins are solitary animals with relatively small population sizes, reflecting their endangered status^10^. However, it cannot be excluded that pangolins acquired their 2019-nCoV related viruses independently from bats or another animal host. It is also notable that both lineages of pangolin coronaviruses were obtained from trafficked Malayan pangolins, likely originating from Southeast Asia, and there is a marked lack of knowledge of the viral diversity maintained by this animal in regions where it is indigenous. Undoubtedly, the extent of virus transmission in pangolin populations requires additional investigation, but the repeated occurrence of infections with 2019-nCoV related coronaviruses in Guangxi and Guangdong provinces suggests that this animal may be a potentially important host in coronavirus emergence.

Coronaviruses, including those related to 2019-nCoV, are clearly present in many wild mammals in Asia^5,6,7,11^. Although the epidemiology, pathogenicity, interspecies infectivity and transmissibility of coronaviruses in pangolins remains to be studied, the data presented here strongly suggests that handling these animals requires considerable caution, and that their sale in wet markets should be strictly prohibited. Further surveillance on pangolins in the natural environment in China and Southeast Asia are clearly needed to understand their role in the emergence of 2019-nCoV and the risk of future zoonotic transmission.

## Methods

### Ethics Statement

The animals studied here were rescued and treated by the Guangxi Zhuang Autonomous Region Terrestrial Wildlife Medical-aid and Monitoring Epidemic Diseases Research Center under the ethics approval (wild animal treatment regulation No. [2011] 85). The samples were collected following the procedure guideline (Pangolins Rescue Procedure, November, 2016).

### Sample collection, viral detection and sequencing of pangolins in Guangxi

We received frozen tissue samples of 18 pangolins (*Manis javanica*) from Guangxi Medical University, China, which were collected between August 2017 – January 2018. These pangolins were seized by the Guangxi Customs during their routine anti-smuggling operations. All animal individuals comprised samples from multiple organs including lungs, intestine and blood, with the exception of six individuals for which only lung tissues were available, five with mixed intestine and lung tissues only, one with intestine tissues only, and one comprising two blood samples. Using the intestine-lung mixed sample we were able to isolate a novel *Betacoronavirus* using the Vero-E6 cell line (Extended Data Figure S1). A High Pure Viral RNA Kit (Roche, Switzerland) was used for RNA extraction on all 43 samples. For RNA sequencing, a sequencing library was constructed using an Ion Total RNA-Seq Kit v2 (Thermo Fisher Scientific, MA, USA), and the library was subsequently sequenced using an Ion Torrent S5 sequencer (Thermo Fisher Scientific). Reverse Transcription was performed using an SuperScript III First-Strand Synthesis System for RT-PCR (Thermo Fisher Scientific, MA, USA). DNA libraries were constructed using the NEBNext Ultra II DNA Library Prep Kit and sequenced on a MiSeq sequencer. The NGS QC Toolkit V2.3.3 was used to remove low-quality and short reads. Both BLASTn and BLASTx were used to search against a local virus database, utilizing the data available at NCBI/GenBank. Genome sequences were assembled using the CLC Genomic Workbench v.9.0. To fill gaps in high throughput sequencing and obtain the whole viral genome sequence, amplicon primers based on the bat SARS-like coronavirus ZC45 (GenBank accession number MG772933) sequence were designed for amplicon-based sequencing.

A total of six samples (including the virus isolate) contained reads that matched a *Betacoronavirus* (Extended Data Table S1). We obtained near complete genomes from these samples (98%, compared to 2019-nCoV), with the virus genomes denoted as GX/P1E, GX/P2V, GX/P3B, GX/P4L, GX/P5E and GX/P5L. Based on these genome sequences, we designed primers for qPCR to confirm the positivity of the original tissue samples (Extended Data Table S4). This revealed an original lung tissue sample that was also qPCR positive, in addition to the six original samples with coronavirus reads. We further tested an addition 19 samples (nine intestine tissues, ten lung tissues), from 12 smuggled pangolins sampled between May-July 2018 by the group from Guangxi Medical University. The genome sequences of GX/P1E, GX/P2V, GX/P3B, GX/P4L, GX/P5E and GX/P5L were submitted to GenBank and GISAID databases, their accession numbers will be available as soon as it is generated.

### Sample collection, viral detection and sequencing of pangolins in Guangdong

After the start of the 2019-nCoV outbreak, the Guangzhou Customs Technology Center re-examined their five archived pangolin samples (two skin swabs, one unknown tissue, one scale) obtained in anti-smuggling operations undertaken in March 2019. RNA was extracted from all five samples (Qiagen, USA), and was subjected to high-throughput RNA sequencing on the Illumina HiSeq platform by Vision medicals, Guangdong, China. The scale sample was found to contain coronavirus reads using BLAST methods. These reads were quality assessed, cleaned and assembled into contigs by both *de novo* (MEGAHIT v1.1.3^12^) and using reference (BWA v0.7.13^13^) assembly methods, with BetaCoV/Wuhan/WIV04/2019 as reference. The contigs were combined, and approximately 72% of the coronavirus genome (21,505bp) was obtained. This sequence was denoted as pangolin CoV GD/P2S.

Liu *et al*. recently published a meta-transcriptomic study of pangolins^7^ and deposited 21 RNA-seq raw files on the SRA database (https://www.ncbi.nlm.nih.gov/sra). We screened these raw read files using BLAST methods and found that five (SRR10168374, SRR10168376, SRR10168377, SRR10168378 and SRR10168392) contained reads that mapped to 2019-nCoV. These reads were subjected to quality assessment, cleaning and then *de novo* assembly using MEGAHIT^12^ and reference assembly using BWA^13^. These reads were then merged and curated in a pileup alignment file to obtain the consensus sequences. This combined consensus sequence is 25,753bp in length (about 86.3% of BetaCoV/Wuhan/WIV04/2019) and denoted pangolin CoV GD/P1L. Notably, it has 66.8% overlap and only 0.21% divergence (i.e. a sequence identify of 99.79%) with the GD/P2S sequence. Since their genetic difference is so low, for the recombination analysis we merged the GD/P1L and GD/P2S sequences into a single consensus sequence to minimize gap regions within any sequences.

The viral genome organizations of Guangxi and Guangdong pangolin coronaviruses were similar to 2019-nCoV. They had a total number of 9 open reading frames (ORFs) and shared the same gene order of ORF1ab replicase, envelope glycoprotein spike (S), envelope (E), membrane (M), nucleocapsid (N), plus other predicted ORFs. Detailed comparison of the ORF length and similarity with 2019-nCoV and bat coronavirus RaTG13 is provided in Extended Table S5.

### Sequence, phylogenetic and recombination analyses

The human 2019-nCoV and bat RaTG13 coronavirus genome sequences were downloaded from Virological.org (http://virological.org) and the GISAID (https://www.gisaid.org) databases on 17 January 2020, with the data kindly shared by the submitters (Extended Data Table S2). Other coronaviruses (subgenus *Sarbecovirus*) were downloaded from GenBank (Extended Data Table S3) and compared to those obtained here. We constructed a multiple sequence alignment of their complete genomes and individual genes using MAFFT v7.273^14^. Maximum likelihood phylogenies were estimated using PhyML v3.1^15^, utilizing the GTR+I+Γ model of nucleotide substitution with 1,000 bootstrap replicates. To investigate potential recombination events, we implemented a window sliding approach to determine the changing patterns of sequence similarity and phylogenetic clustering between the query and the reference sequences, as well as a scanning of phylogenetic clusters performed directly from the multiple sequence alignment. Maximum likelihood trees were estimated from each window extraction (i.e. genome regions 1 to 8) using PhyML as described above.

## Acknowledgements

We thank Prof. Wu-Chun Cao, Dr. Na Jia, Dr. Ya-Wei Zhang, Dr. Jia-Fu Jiang, Dr. Bao-Gui Jiang, and their team in State Key Laboratory of Pathogen and Biosecurity, Beijing Institute of Microbiology and Epidemiology, Beijing for their substantial contributions to this study, including coordinating among research parties, conducting virus isolation, qPCR and sequencing. We also thank Guangzhou Customs Technology Center, Guangzhou for re-examining their archived pangolin samples and providing the data. All the above persons agree on the content of the study, authorship arrangement and publication. We thank the staff of the Guangxi and Guangdong Custom Bureau for their laborious anti-smuggling operations. We thank all the scientists who kindly shared their genomic sequences of the coronaviruses used in this study.

This work was supported by research grants from National Key Plan for Scientific Research and Development of China (2016YFD0500302; 2017YFE0190800), funding for Guangdong-Hongkong-Macau Joint Laboratory (2019B121205009), The National Natural Science Foundation of China (NSFC) Excellent Young Scientists Fund (Hong Kong and Macau) (31922087), Li Ka Shing Foundation, National Institutes of Health (HHSN272201400006C), and the Australian Research Council (FL170100022).

## Author contributions

Y.G. and Y.L.H. designed and supervised research. W.W., W.J.L. and L.F.L. collected samples and conducted genome sequencing. M.H.H.S, X.B.N. and T.T.Y.L. performed genome assembly and annotation. Y.G.T., T.T.Y.L., M.H.H.S, X.B.N, W.Y.M.C. and Y.S.L. performed genome analysis and interpretation. T.T.Y.L. and E.C.H.wrote the paper. H.C.Z., Y.L.H., G.M.L and Y.G. joined the data interpretation and edited the paper.

## Materials & correspondence

Correspondence and requests for materials should be addressed to Prof. Yi Guan

## Data availability

Data that support the findings of this study have been deposited in GenBank database with the xxxx-xxxx accession codes and SRA database with xxxx-xxxx accession codes.

## Extended Data

**Figure S1.**
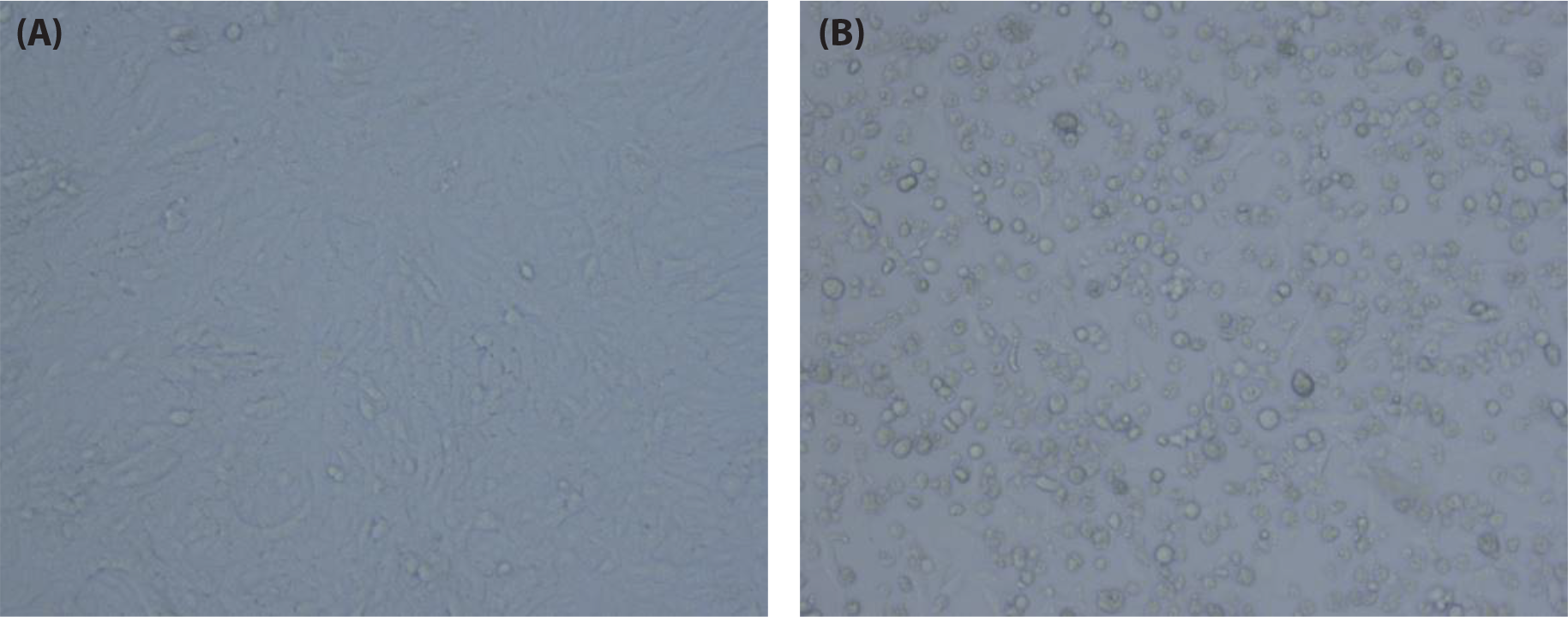
Microscopic image of the cytopathic effect in virus isolation using Vero E6. (A) Negative control of Vero E6 cell line. (B) Cytopathic effect seen in viral culture (5 days post inoculation).

**Table S1.** High-throughput sequencing results of the samples with coronavirus reads

| Source location | Animal | Sample type | Sample number | Sequencing raw data ID |
| --- | --- | --- | --- | --- |
| Guangxi | Pangolin | Intestine | GX/P1E | Data submission in process; Identifier will be available as soon as it is generated. |
| Guangxi | Pangolin | Virus isolate from intestine-lung mixed samples | GX/P2V | Data submission in process; Identifier will be available as soon as it is generated. |
| Guangxi | Pangolin | Blood | GX/P3B | Data submission in process; Identifier will be available as soon as it is generated. |
| Guangxi | Pangolin | Lung | GX/P4L | Data submission in process; Identifier will be available as soon as it is generated. |
| Guangxi | Pangolin | Intestine | GX/P5E | Data submission in process; Identifier will be available as soon as it is generated. |
| Guangxi | Pangolin | Lung | GX/P5L | Data submission in process; Identifier will be available as soon as it is generated. |
| Guangdong | Pangolin | Scale | GD/P2S | Data submission in process; Identifier will be available as soon as it is generated. |

**Table S2.** Acknowledgement of sharing of 2019-nCoV genome sequences from the Virological.org and the GISAID databases. We gratefully thank the authors listed below for sharing their genomic sequences of coronaviruses analyzed in this study.

| Accession ID | Virus name | Location | Collection date | Originating lab | Submitting lab | Authors |
| --- | --- | --- | --- | --- | --- | --- |
| Virological.org sequence (NC_045512.2) | BetaCoV/Wuhan-Hu-1/2019 | China / Wuhan | 2019-12 | National Institute for Communicable Disease Control and Prevention (ICDC) Chinese Center for Disease Control and Prevention (China CDC) | National Institute for Communicable Disease Control and Prevention (ICDC) Chinese Center for Disease Control and Prevention (China CDC) | Zhang,Y.-Z., Wu,F., Chen,Y.-M., Pei,Y.-Y., Xu,L., Wang,W., Zhao,S., Yu,B., Hu,Y., Tao,Z.-W., Song,Z.-G., Tian,J.-H., Zhang,Y.-L., Liu,Y., Zheng,J.-J., Dai,F.-H., Wang,Q.-M., She,J.-L. and Zhu,T.-Y. |
| EPI_ISL_402131 | BetaCoV/bat/Yunnan/RaTG13/2013 | China / Yunnan Province / Pu'er City | 2013-07-24 | Wuhan Institute of Virology, Chinese Academy of Sciences | Wuhan Institute of Virology, Chinese Academy of Sciences | Yan Zhu, Ping Yu, Bei Li, Ben Hu, Hao-Rui Si, Xing-Lou Yang, Peng Zhou, Zheng-Li Shi |
| EPI_ISL_402121 | BetaCoV/Wuhan/IVDC-HB-05/2019 | China / Hubei Province / Wuhan City | 2019-12-30 | National Institute for Viral Disease Control and Prevention, China CDC | National Institute for Viral Disease Control and Prevention, China CDC | Wenjie Tan, Xuejun Ma, Xiang Zhao, Wenling Wang, Yongzhong Jiang, Roujian Lu, Ji Wang, Peihua Niu, Weimin Zhou, Faxian Zhan, Weifeng Shi, Baoying Huang, Jun Liu, Li Zhao, Yao Meng, Fei Ye, Na Zhu, Xiaozhou He, Peipei Liu, Yang Li, Jing Chen, Wenbo Xu, George F. Gao, Guizhen Wu |
| EPI_ISL_402120 | BetaCoV/Wuhan/IVDC-HB-04/2020 | China / Hubei Province / Wuhan City | 2020-01-01 | National Institute for Viral Disease Control and Prevention, China CDC | National Institute for Viral Disease Control and Prevention, China CDC | Wenjie Tan, Xiang Zhao, Wenling Wang, Xuejun Ma, Yongzhong Jiang, Roujian Lu, Ji Wang, Weimin Zhou, Peihua Niu, Peipei Liu, Faxian Zhan, Weifeng Shi, Baoying Huang, Jun Liu, Li Zhao, Yao |
|  |  |  |  |  |  | Meng, Xiaozhou He,<br>Fei Ye, Na Zhu, Yang<br>Li, Jing Chen, Wenbo<br>Xu, George F. Gao,<br>Guizhen Wu |
| EPI_ISL_<br>402124 | BetaCoV/Wuhan/<br>WIV04/2019 | China /<br>Hubei<br>Province<br>/ Wuhan<br>City | 2019-12-30 | Wuhan Jinyintan<br>Hospital | Wuhan Institute<br>of Virology,<br>Chinese Academy<br>of Sciences | Peng Zhou, Xing-Lou<br>Yang, Ding-Yu Zhang,<br>Lei Zhang, Yan Zhu,<br>Hao-Rui Si, Zhengli Shi |
| EPI_ISL_<br>402123 | BetaCoV/Wuhan<br>/IPBCAMS-WH-01/2019 | China /<br>Hubei<br>Province<br>/ Wuhan<br>City | 2019-12-24 | Institute of<br>Pathogen Biology,<br>Chinese Academy<br>of Medical<br>Sciences & Peking<br>Union Medical<br>College | Institute of<br>Pathogen<br>Biology, Chinese<br>Academy of<br>Medical Sciences<br>& Peking Union<br>Medical College | Lili Ren, Jianwei Wang,<br>Qi Jin, Zichun Xiang,<br>Zhiqiang Wu, Chao Wu,<br>Yiwei Liu |

**Table S3.**
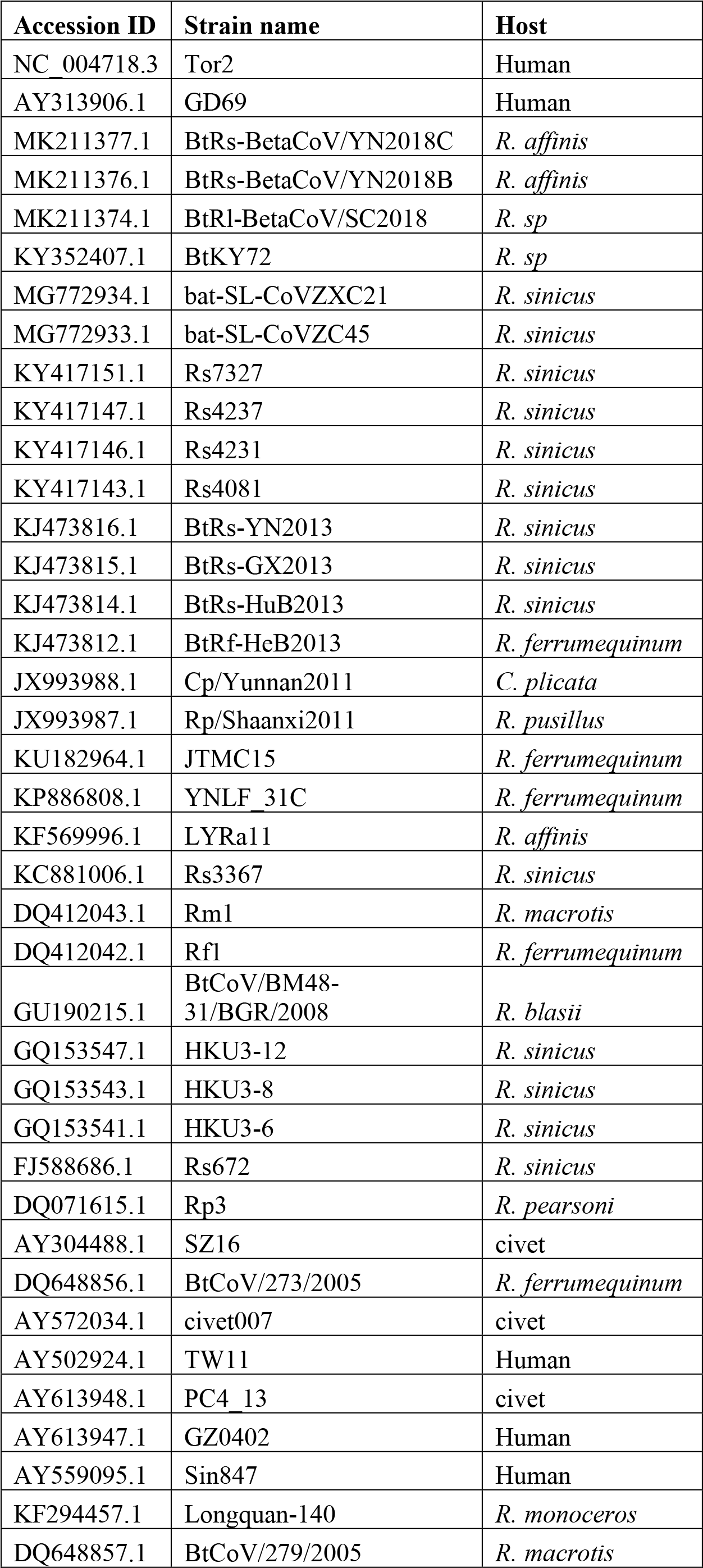
GenBank accession numbers of coronavirus sequences used in this study.

| Accession ID | Strain name | Host |
| --- | --- | --- |
| NC_004718.3 | Tor2 | Human |
| AY313906.1 | GD69 | Human |
| MK211377.1 | BtRs-BetaCoV/YN2018C | *R. affinis* |
| MK211376.1 | BtRs-BetaCoV/YN2018B | *R. affinis* |
| MK211374.1 | BtRl-BetaCoV/SC2018 | *R. sp* |
| KY352407.1 | BtKY72 | *R. sp* |
| MG772934.1 | bat-SL-CoVZXC21 | *R. sinicus* |
| MG772933.1 | bat-SL-CoVZC45 | *R. sinicus* |
| KY417151.1 | Rs7327 | *R. sinicus* |
| KY417147.1 | Rs4237 | *R. sinicus* |
| KY417146.1 | Rs4231 | *R. sinicus* |
| KY417143.1 | Rs4081 | *R. sinicus* |
| KJ473816.1 | BtRs-YN2013 | *R. sinicus* |
| KJ473815.1 | BtRs-GX2013 | *R. sinicus* |
| KJ473814.1 | BtRs-HuB2013 | *R. sinicus* |
| KJ473812.1 | BtRf-HeB2013 | *R. ferrumequinum* |
| JX993988.1 | Cp/Yunnan2011 | *C. plicata* |
| JX993987.1 | Rp/Shaanxi2011 | *R. pusillus* |
| KU182964.1 | JTMC15 | *R. ferrumequinum* |
| KP886808.1 | YNLF_31C | *R. ferrumequinum* |
| KF569996.1 | LYRa11 | *R. affinis* |
| KC881006.1 | Rs3367 | *R. sinicus* |
| DQ412043.1 | Rm1 | *R. macrotis* |
| DQ412042.1 | Rf1 | *R. ferrumequinum* |
| GU190215.1 | BtCoV/BM48-31/BGR/2008 | *R. blasii* |
| GQ153547.1 | HKU3-12 | *R. sinicus* |
| GQ153543.1 | HKU3-8 | *R. sinicus* |
| GQ153541.1 | HKU3-6 | *R. sinicus* |
| FJ588686.1 | Rs672 | *R. sinicus* |
| DQ071615.1 | Rp3 | *R. pearsoni* |
| AY304488.1 | SZ16 | civet |
| DQ648856.1 | BtCoV/273/2005 | *R. ferrumequinum* |
| AY572034.1 | civet007 | civet |
| AY502924.1 | TW11 | Human |
| AY613948.1 | PC4_13 | civet |
| AY613947.1 | GZ0402 | Human |
| AY559095.1 | Sin847 | Human |
| KF294457.1 | Longquan-140 | *R. monoceros* |
| DQ648857.1 | BtCoV/279/2005 | *R. macrotis* |

**Table S4.**
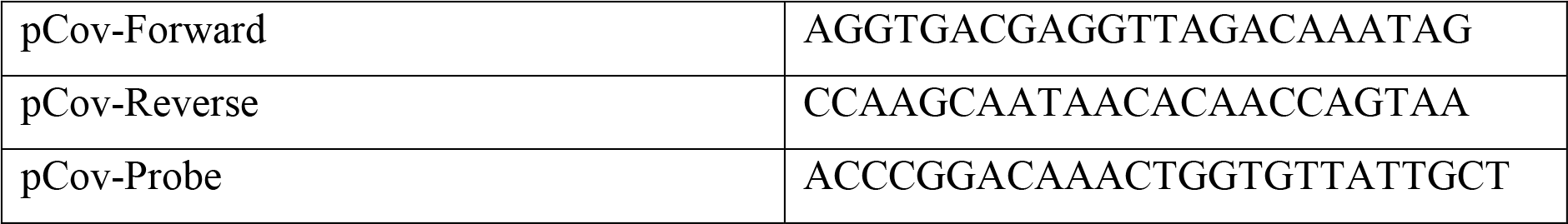
Primers used for qPCR detection of pangolin associated coronavirus

**Table S5.**
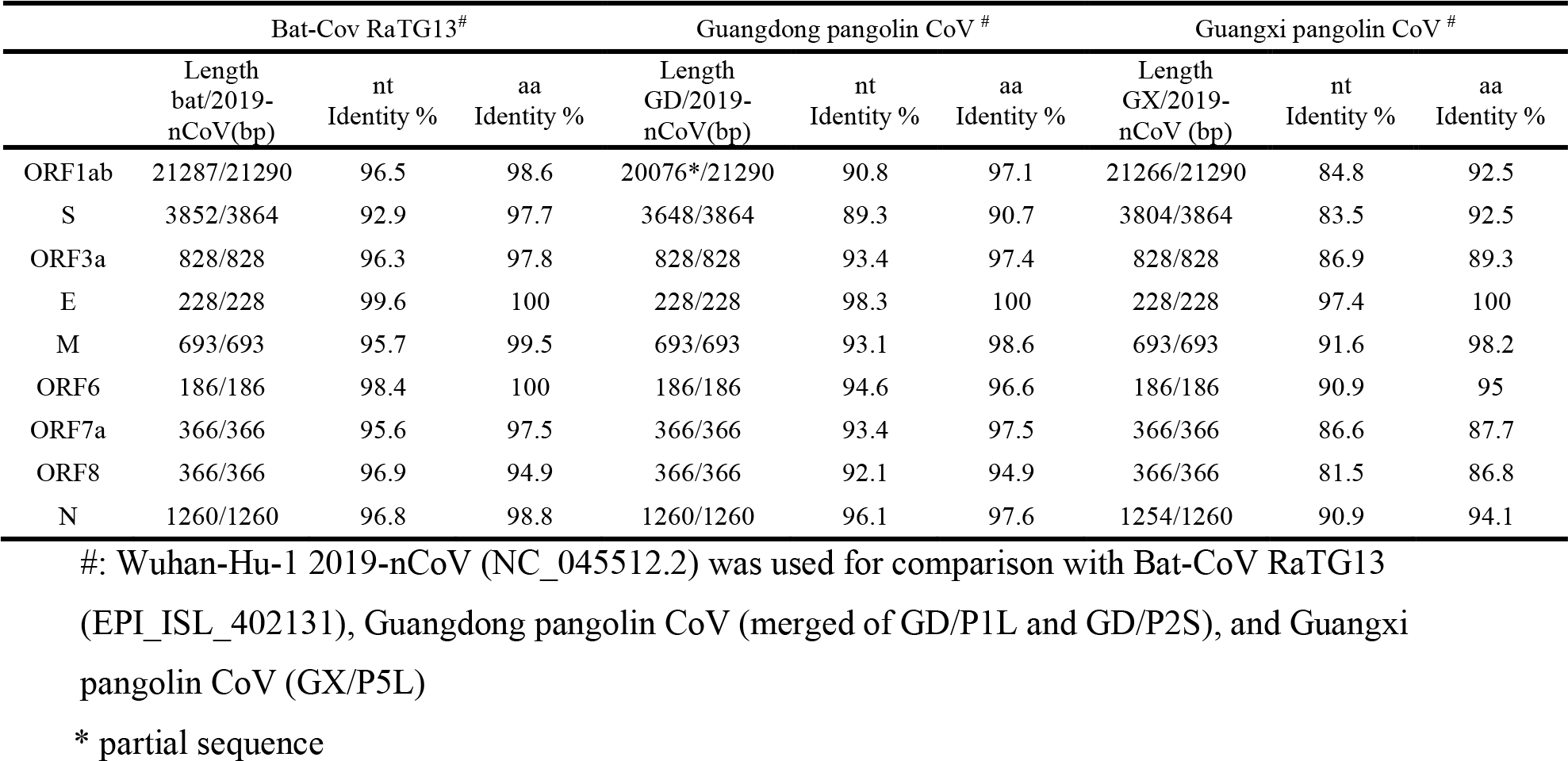
Genomic comparison of 2019-nCoV with Bat-Cov RaTG13, Guangdong pangolin CoV and Guangxi pangolin CoV.

|  | Bat-Cov RaTG13<sup>#</sup> |  |  | Guangdong pangolin CoV <sup>#</sup> |  |  | Guangxi pangolin CoV <sup>#</sup> |  |  |
| --- | --- | --- | --- | --- | --- | --- | --- | --- | --- |
|  | Length<br>bat/2019-nCoV(bp) | nt<br>Identity % | aa<br>Identity % | Length<br>GD/2019-nCoV(bp) | nt<br>Identity % | aa<br>Identity % | Length<br>GX/2019-nCoV (bp) | nt<br>Identity % | aa<br>Identity % |
| ORF1ab | 21287/21290 | 96.5 | 98.6 | 20076*/21290 | 90.8 | 97.1 | 21266/21290 | 84.8 | 92.5 |
| S | 3852/3864 | 92.9 | 97.7 | 3648/3864 | 89.3 | 90.7 | 3804/3864 | 83.5 | 92.5 |
| ORF3a | 828/828 | 96.3 | 97.8 | 828/828 | 93.4 | 97.4 | 828/828 | 86.9 | 89.3 |
| E | 228/228 | 99.6 | 100 | 228/228 | 98.3 | 100 | 228/228 | 97.4 | 100 |
| M | 693/693 | 95.7 | 99.5 | 693/693 | 93.1 | 98.6 | 693/693 | 91.6 | 98.2 |
| ORF6 | 186/186 | 98.4 | 100 | 186/186 | 94.6 | 96.6 | 186/186 | 90.9 | 95 |
| ORF7a | 366/366 | 95.6 | 97.5 | 366/366 | 93.4 | 97.5 | 366/366 | 86.6 | 87.7 |
| ORF8 | 366/366 | 96.9 | 94.9 | 366/366 | 92.1 | 94.9 | 366/366 | 81.5 | 86.8 |
| N | 1260/1260 | 96.8 | 98.8 | 1260/1260 | 96.1 | 97.6 | 1254/1260 | 90.9 | 94.1 |

<sup>#</sup>: Wuhan-Hu-1 2019-nCoV (NC\_045512.2) was used for comparison with Bat-CoV RaTG13
(EPI\_ISL\_402131), Guangdong pangolin CoV (merged of GD/P1L and GD/P2S), and Guangxi pangolin CoV (GX/P5L)
\* partial sequence

